# Estimation of Jacquard’s genetic identity coefficients with bi-allelic variants by constrained least-squares

**DOI:** 10.1101/2024.03.25.586682

**Authors:** Jan Graffelman, Bruce S. Weir, Jérôme Goudet

## Abstract

The Jacquard genetic identity coefficients are of fundamental importance in relatedness research. We address the estimation of these coefficients as well as other relatedness parameters that derive from them such as kinship and inbreeding coefficients using a concise matrix framework. Estimation of the Jacquard coefficients via likelihood methods and the expectation–maximization algorithm is computationally very demanding for large numbers of polymorphisms. We propose a constrained least squares approach to estimate the Jacquard coefficients. A simulation study shows constrained least squares achieves root-mean-squared errors that are comparable with those of the maximum likelihood approach, in particular when founder allele frequencies are unknown, while obtaining enormous computational savings.

## 1 Introduction

The estimation of the degree of genetic relatedness, either by using pedigrees or molecular marker data is of keen interest for a variety of purposes (Weir et al. 2006). It is, among others, useful for establishing genealogies, for paternity testing, and for maintaining genetic diversity in breeding programmes with endangered species. Accounting for relatedness is crucial in genetic association studies (Astle & Balding 2009). The definition of the concept of alleles that derive from a common ancestor as identical-by-descent (ibd) alleles (Malécot 1970) was foundational for relatedness research. Harris (1964) enumerated 15 modes of identity-by-descent, which reduce to nine modes if the paternal and maternal origins of the alleles are not distinguished. These modes were represented pictorially by Jacquard (1972, 1974), and are nowadays commonly referred to as Jacquard’s coefficients. Figure 1 shows the nine condensed states where blue lines indicate an ibd relationship between two alleles. Vertical lines show ibd relationships between individuals of a pair, horizontal lines refer to ibd relationships within an individual, i.e. these refer to an inbred state. We use the symbol Δ_*i*_ to either refer to the particular mode or its probability. For patterns Δ_1_ and Δ_7_ the two individuals share two ibd alleles among them; for patterns Δ_3_, Δ_5_ and Δ_8_ they share one, and for the remaining states they share none. States Δ_1_ through Δ_6_ all refer to inbred states. When there is no inbreeding, the number of states reduces to three (Δ_7_, Δ_8_ and Δ_9_), and their relative probabilities are known as the Cotterman coefficients (1940). Under non-inbred conditions Thompson (1976) showed the Cotterman coefficients are limited to a subspace of the two-dimensional three-part simplex, satisfying 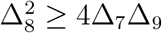.

**Figure 1.**
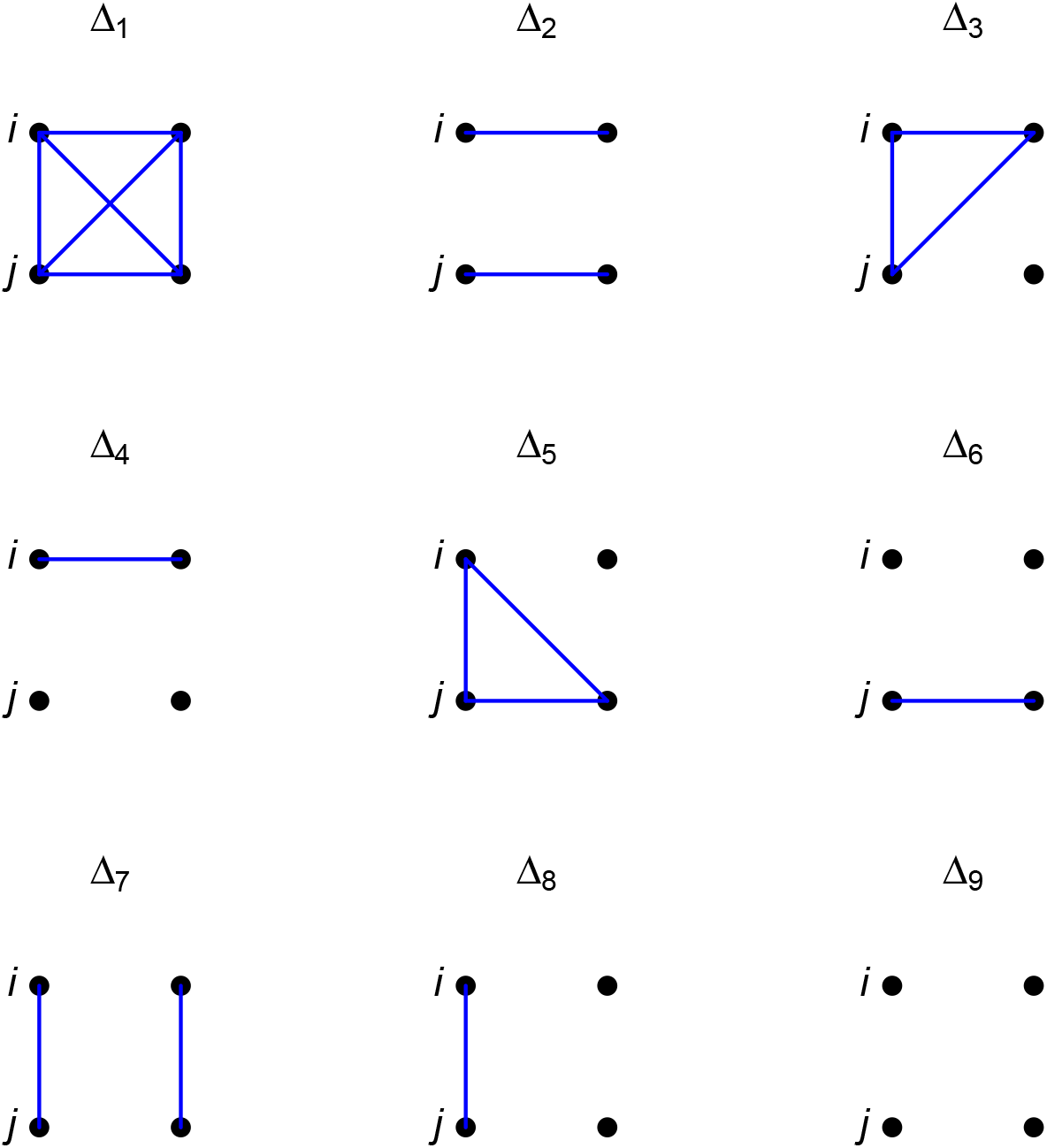
The ibd patterns for a pair (*i, j*) of individuals. Dots represent alleles. Blue lines connect pairs of alleles that are ibd.

Probabilities of observed genotypes for pairs of individuals are readily related to the Jacquard coefficients for given allele frequencies (Cockerham 1971). We will use a 0 and a 1 to respectively represent the major and minor allele at a bi-allelic locus, and use 0/0, 0/1 and 1/1 to represent the corresponding diploid genotypes. Thus, for a bi-allelic variant with minor allele frequency (maf) *p* and major allele frequency *q*, the probability of observing either two minor homozygotes or two major homozygotes is, given state Δ_1_, *p* or *q* respectively. Likewise, state Δ_2_ is compatible only with genotype pairs (0/0,1/1), (0/0,0/0), (1/1,1/1) and (1/1,0/0), which will have probabilities *pq, p*^2^, *q*^2^ and *qp* respectively, since the first allele of an individual is necessarily the same as the second. By the same token, for each given mode the joint genotype probabilities can be developed for all nine possible genotype pairs and are given in Table 1.

**Table 1:** The bi-allelic condensed system, consisting of joint genotype probabilities for given ibd patterns and allele frequencies.

| Nr. | Genotype pair | $\Delta_1$ | $\Delta_2$ | $\Delta_3$ | $\Delta_4$ | $\Delta_5$ | $\Delta_6$ | $\Delta_7$ | $\Delta_8$ | $\Delta_9$ |
| --- | --- | --- | --- | --- | --- | --- | --- | --- | --- | --- |
| 1 | (0/0,0/0) | $q$ | $q^2$ | $q^2$ | $q^3$ | $q^2$ | $q^3$ | $q^2$ | $q^3$ | $q^4$ |
| 2 | (0/0,0/1) | 0 | 0 | $pq$ | $2pq^2$ | 0 | 0 | 0 | $pq^2$ | $2pq^3$ |
| 3 | (0/0,1/1) | 0 | $pq$ | 0 | $p^2q$ | 0 | $pq^2$ | 0 | 0 | $p^2q^2$ |
| 4 | (0/1,0/0) | 0 | 0 | 0 | 0 | $pq$ | $2pq^2$ | 0 | $pq^2$ | $2pq^3$ |
| 5 | (0/1,0/1) | 0 | 0 | 0 | 0 | 0 | 0 | $2pq$ | $p^2q + pq^2$ | $4p^2q^2$ |
| 6 | (0/1,1/1) | 0 | 0 | 0 | 0 | $pq$ | $2p^2q$ | 0 | $p^2q$ | $2p^3q$ |
| 7 | (1/1,0/0) | 0 | $pq$ | 0 | $pq^2$ | 0 | $p^2q$ | 0 | 0 | $p^2q^2$ |
| 8 | (1/1,0/1) | 0 | 0 | $pq$ | $2p^2q$ | 0 | 0 | 0 | $p^2q$ | $2p^3q$ |
| 9 | (1/1,1/1) | $p$ | $p^2$ | $p^2$ | $p^3$ | $p^2$ | $p^3$ | $p^2$ | $p^3$ | $p^4$ |

Table 1 has been published in different forms by several authors (Cockerham 1971, Weir et al. 2006, Anderson & Weir 2007, Csűrös 2014, Laporte et al. 2017, Wang 2022), depending on whether or not two or more alleles are considered, and depending on whether the order of the individuals in a pair is taken into account or not. Anderson & Weir (2007) developed a (multi-allelic) extended parametrization of Table 1 in order to allow for population sub-structure, i.e., allowing for a subpopulation whose allele frequency has drifted away from the original parent population. As given here, Table 1 is strictly for the bi-allelic case, the order of the alleles of an individual is considered irrelevant (i.e., heterozygotes 0/1 and 1/0 are not distinguished), but the order of the individuals in a pair is taken into account. A homogeneous population with no differentiation of allele frequencies is assumed throughout.

If we define **g** as the 9 *×* 1 vector of (marginal) joint genotype probabilities, then by the law of total probability, we have

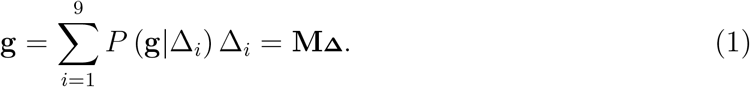

where **M** is the 9 *×* 9 matrix given in Table 1, whose entries are determined by a single parameter, the minor allele frequency, and **Δ** = (Δ_1_, Δ_2_, …, Δ_9_)^*′*^ the column vector containing the coefficients for one particular pair (*i,j*) of individuals. Alternative parametrizations of the system in terms of genetic correlations coefficients (Ackerman et al. 2017) are possible, but not considered here. Equation (1) refers to the so-called condensed coefficients (Jacquard 1974), and we will refer to it as the *bi-allelic condensed system*.

Up to only a few years ago, much of relatedness research mostly focused on the estimation of the Cotterman coefficients and the derived kinship coefficient, where the latter is defined as the probability that two alleles, each taken at random from an individual, are ibd.

The estimation of the ibd coefficients by maximum likelihood (Thompson 1975), assuming non-inbred individuals and known allele frequencies, was a milestone achievement for relatedness research. Under absence of inbreeding, the kinship coefficient is obtained from the Cotterman coefficients as 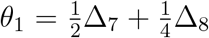. Milligan (2003) performed an extensive study comparing the different estimators of the kinship coefficient, and generally recommended the maximum likelihood estimator across a broad spectrum of conditions. Hitherto, very few papers have addressed the estimation of the full set of Jacquard coefficients.

Nowadays, the ever-growing amount of available genetic information and computational resources have lead to an increased interest in the estimation of relatedness parameters such as coancestry, individual inbreeding coefficients, and others. Multiple estimators have recently been proposed for these parameters, including allele-sharing estimators (Weir & Goudet 2017, Goudet et al. 2018) for coancestry and inbreeding that account for genetic sampling (Weir 1996). Most relatedness parameters of interest can be derived from Jacquard’s coefficients, and correspondingly some attempts have been made to estimate the full set of nine condensed Jacquard coefficients, using different methods (Hanghøj et al. 2019, Laporte et al. 2017, Wang 2022), despite the fact that for bi-allelic data, Jacquard’s coefficients have been shown not to be identifiable (Csűrös 2014).

In this article we propose to estimate the nine Jacquard coefficients and derived quantities by using a constrained least squares (cls) criterion, as described in Section 2 below. We perform pedigree-based simulations to compare cls and em estimates, and also address the computational cost of scaling up the algorithms to a full genome.

## 2 Theory

### Notation

We first develop our notation. Since many of the quantities involved such as Jacquard coefficients, kinship coefficients and others are pairwise quantities, we have found it convenient to use matrix notation, using bold lowercase and uppercase letters to indicate vectors and matrices respectively. We use a 1 to denote the minor allele of a SNP, and 0 to denote its major allele. We use *p*_*i*_ to refer to minor allele frequency of the *i*th SNP and *q*_*i*_ = 1 − *p*_*i*_ for its major allele frequency. We use, following Jacquard (1974), the scalar Δ_*k*_ to denote the probability of state *k* for pair of individuals (*i, j*). For convenience, we store all pairwise coefficients in *n × n* subindexed matrices **Δ**_1_, **Δ**_2_, … **Δ**_9_, e.g., the element in row *i* and column *j* of matrix **Δ**_1_ contains the probability Δ_1_ for the pair of individuals (*i, j*). When convenient, we will make use of vector **Δ** (small case without subscript), and **Δ** = (Δ_1_, …, Δ_9_) refers to the set of nine Jacquard coefficients for a particular pair (*i, j*). We start by noting that the probabilities of the nine possible states constitute a closed simplex (Thompson 1978), given by

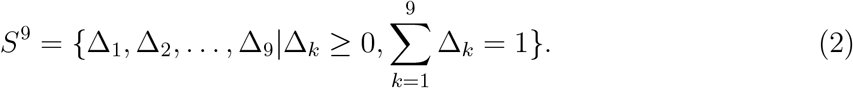

We enumerate some of the well-known quantities that are derived from the Jacquard coefficients. The coancestry or kinship coefficient is given by

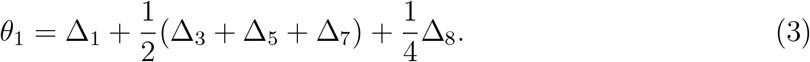

The inbreeding coefficients of the *i*th and *j*th individual are given by

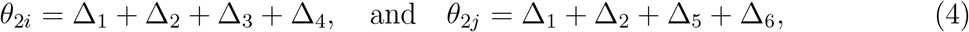

where we use subscripts 2*i* and 2*j* to refer to individual inbreeding; these quantities have been termed *θ*_2*A*_, *θ*_2*B*_ or *f*_*A*_, *f*_*B*_ by others (Csűros 2014, Jacquard 1974). The probability of at least one pair of ibd alleles among three randomly selected alleles, of the four carried by individuals (*i, j*), is given by

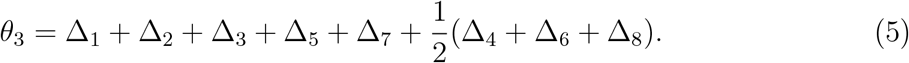

finally, we define

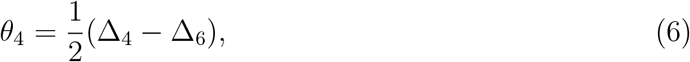

making for five identifiable relatedness parameters (Csűrös 2014). A statistical model is identifiable if there is a one-to-one correspondence between the values of the parameters of the model and the probability distribution of the data. For bi-allelic polymorphisms, the set of condensed Jacquard coefficients is not identifiable because two different sets of coefficients can generate the same probability distribution of joint genotypes, as illustrated in Appendix A. All five identifiable relatedness parameters above of a pair can be conveniently obtained by a linear transformation (**Q**, of rank five) of the Jacquard coefficients as ***θ*** = **Q**_**Δ**_, i.e.

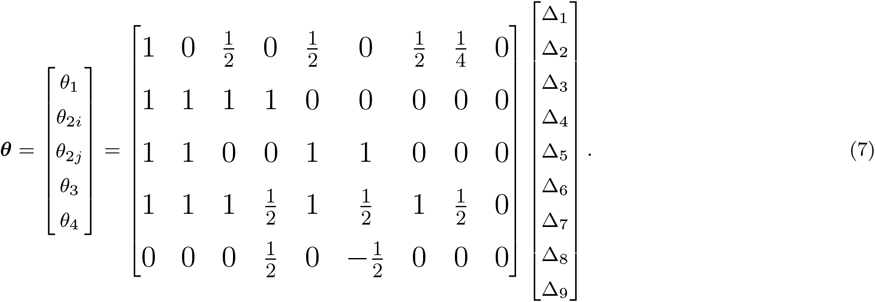

We note that the vector of identifiable relatedness parameters ***θ*** is not unique, and that alternative vectors of identifiable relatedness parameters can be obtained by defining linear combinations of the rows of **Q** (Csűrös 2014, Theorems 4 and 5). It is insightful to further develop the matrix notation, and the simplex property implies that

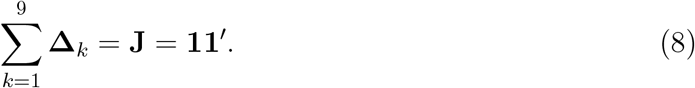

Note that for states Δ_3_ and Δ_5_, an interchange of the two individuals *i* and *j* implies a change from state Δ_3_ to Δ_5_ for one individual, and a change from state Δ_5_ to Δ_3_ for the other (see Figure 1). Correspondingly, **Δ**_3_ and **Δ**_5_ are not symmetric but are each other’s mutual transpose. The same holds true for states Δ_4_ and Δ_6_. For all other states, an interchange of individuals does not bring about a change of state, and we therefore have that

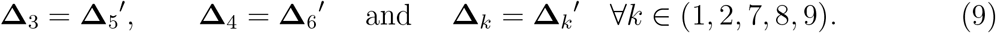

When a Jacquard coefficient of an individual with himself is considered, all states have probability zero except Δ_1_ and Δ_7_, because an individual always shares two ibd alleles with himself, which are either inbred (Δ_1_) or not (Δ_7_). Consequently, we have

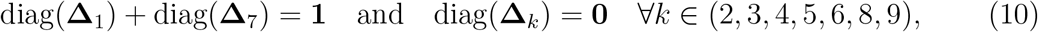

such that only **Δ**_1_ and **Δ**_7_ can have non-zero diagonals, and where operator diag(*·*) extracts the diagonal of a matrix into a column vector. We next develop matrices for relatedness coefficients. The kinship matrix is defined as

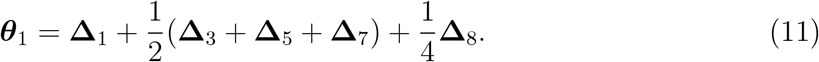

This matrix is symmetric because

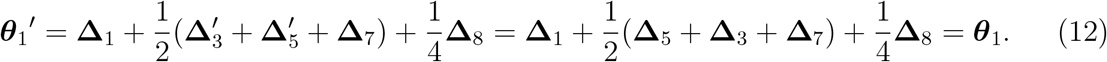

Note that for self-kinship

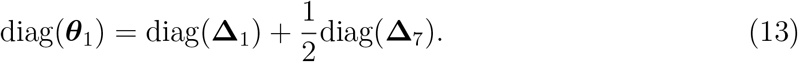

Matrices of inbreeding coefficients are, according to Equation (4), obtained as

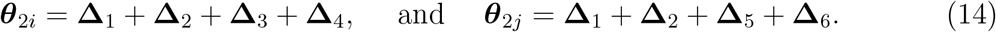

So that

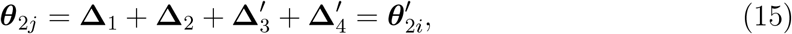

which implies

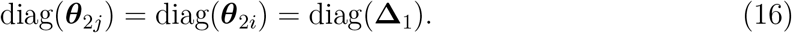

Multiplying (13) by two and combining with (9)

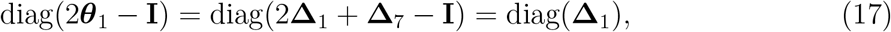

which can be rewritten as

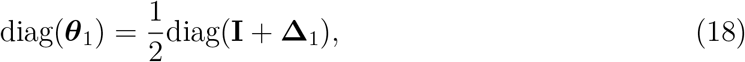

where the latter equation is the matrix formulation of the well-known result that self-kinship relates to inbreeding 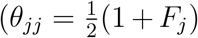, in a fairly usual scalar notation). Inbreeding coefficients for each individual are thus obtained as the diagonal elements of **Δ**_1_, or equivalently, as the row means of ***θ***_2*i*_ or the column means of ***θ***_2*j*_. The obvious matrix formulations for *θ*_3_ and *θ*_4_ are

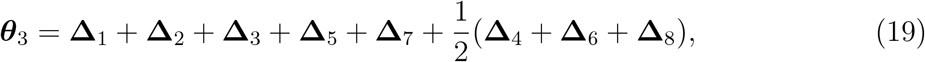

which has diag(***θ***_3_) = **1**, and is symmetric. Finally,

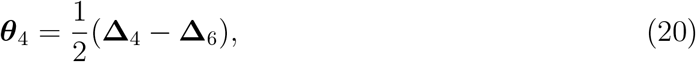

is skew-symmetric 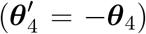. The aforementioned close relationship between states (Δ_3_, Δ_5_) and (Δ_4_, Δ_6_), suggests these states might be joined by summation, reducing the number of parameters to be estimated to seven. This reduction is developed in Appendix B.

Following Thompson (2013), Weir and Goudet (2017) emphasized the relative nature of coancestry and inbreeding, defining the compound quantities of *relative* coancestry and *relative* inbreeding, which we will indicate with *ϑ*_1_ and *ϑ*_2_ respectively. These quantities are readily obtained from the previous expressions. We define the theoretical average coancestry over all *n*(*n* − 1) pairs as

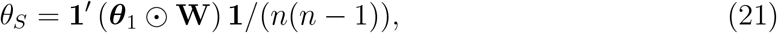

where ⊙ represents the Hadamard product (i.e. elementwise multiplication), and **W** a weight matrix of ones with zeros on the diagonal (**W** = **J** − **I**). The symmetric matrix of relative coancestry coefficients ***ϑ***_1_, is obtained as

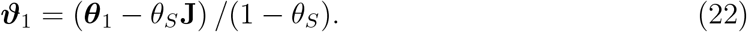

We note that ***ϑ***_1_ precisely contains the relative individual inbreeding coefficients on its diagonal. Let ***ϑ***_2_ be a column vector containing these coefficients. Using Eq. (16) We have that

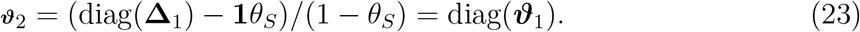

In the remainder, we will use 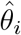 to refer to estimators of the relatedness parameters, and 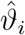 to refer to estimators of the corresponding relative parameters.

### Estimation

Equation (1) describes a theoretical population-genetic model, giving an expected relationship between the joint pairwise genotype probabilities, allele frequencies and Jacquard’s coefficients, obtained under the random mating assumption. Equation (1) has been solved with maximum likelihood procedures, by assuming known allele frequencies and building the multinomial likelihood function by multiplying this equation over loci (Laporte et al. 2017). The latter authors estimate the Jacquard coefficients by ml using an EM algorithm, using the crossing design to improve identifiability of the coefficients.

In this article, we elaborate on the alternative approach initiated by Csűrös (2014) and regard Equation (1) as a system of linear equations that could be solved for **Δ** for each pair (*i, j*) if **g** and **M** were known; one thus would need to estimate both **g** and **M** from the genotype data, prior to estimating **Δ**. Csűrös (2014) pointed out matrix **M** is structurally singular, and is expected to be of rank seven. It is subject to two linear constraints, both for the columns and the rows. When considering the rows of **M**, it is straightforward to show that these constraints amount to **1**^*′*^**M** = **1**^*′*^ and **a**^*′*^**M** = **0**^*′*^, with **a**^*′*^ = (0, 0, 1, −1, 0, 2, −2, 1, −1). Equation (1), viewed as a system of linear equations, can be either consistent or inconsistent, and we address both situations below.

### The consistent system

Equation (1) will constitute a consistent system with infinitely many solutions provided that **M** and **g** are parametrized by exactly the same minor allele probability *p*, and satisfy **a**^*′*^**g** = **a**^*′*^**M**_**Δ**_ = 0. In that case, the system can be solved for *some* particular solution either using Gaussian elimination or by the use of a generalized inverse, such as the Moore-Penrose inverse (Searle 1982). Gaussian elimination will reduce **M** to row-echelon form with trailing rows of zeros, and retains the column-sum-one property. Consequently, the obtained Jacquard coefficients will sum to one, but they can be negative. In most cases, **M** will have rank seven due to the two linear restrictions identified by Csűrös (2014). More precisely, the rank of **M** depends on *p* and is at most seven; e.g. **M** will have rank five whenever *p* = *q* = 0.5. Whenever **M** has rank seven, Gaussian elimination leads to 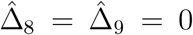, whereas other coefficients can be negative. This may, at first sight, be surprising, for Δ_8_ and Δ_9_ typically correspond to the largest Jacquard coefficients found in practice. However, since the order of the variables in the system of equations is arbitrary, Jacquard coefficients can be set to zero at will by permuting the columns of **M** together with their corresponding elements of **Δ**, clearly showing the coefficients are not identified. A consistent linear system with a structurally singular coefficient matrix can also be resolved using the Moore-Penrose inverse (**M**^+^) of **M**, and estimating **Δ** as 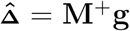. This also gives some particular solution with Jacquard coefficients that sum one, but some of them can be negative. The obvious freedom of the coefficients has been parametrized by Csűrös (2014), and his parametrization can be used to map the coefficients to a set of strictly non-negative Jacquard coefficients (i.e. probabilities) with the transformation

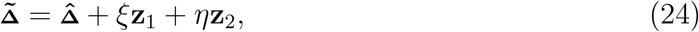

where **z**_1_ = (0, 1, 0, −1, 0, −1, −1, 2, 0) and **z**_2_ = (0, 0, 0, 0, 0, 0, 1, 2, 1)+*pq*(−1, −1, 2, 0, 2, 0, −2, 0, 0) and where *ξ* and *η* are real parameters that are constrained by a set of inequal-ities (Csűrös 2014, Eq. (8)) that warrant the non-negativity of 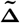; typically this transformation will also render 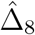 and 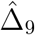 non-zero if one attempted to solve the system by Gaussian elimination. Csűrös (2014) used this result to explore the range of variation of the Jacquard coefficients for an empirical pedigree. Here we use this parametrization with molecular marker data to assess the range of variation of the Jacquard coefficients in that setting. An example case is described in Appendix A. However, it should be recognized that observed joint genotype probabilities generally do not conform to Equation (1); in particular the condition **a**^*′*^**g** = 0 is generally not met. For empirical data, Equation (1) is therefore generally inconsistent, and we therefore capitalize on the inconsistent case developed below.

### The inconsistent system

For given allele frequencies, model (1) can be estimated by averaging, in the unweighted sense, over *L* SNPs such that we need to resolve

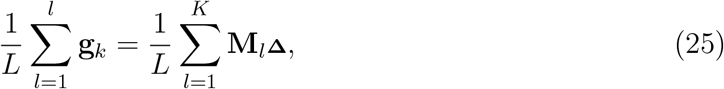

which we write concisely as 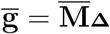, for **Δ**. In general, this system will be inconsistent, but a best-fitting estimate for **Δ** can be found using a least-squares criterion. Retaining a probabilistic interpretation of the Jacquard coefficients, we minimize the residual sum-of-squares given by

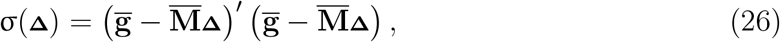

under the restrictions 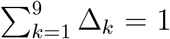 and Δ_*k*_ ≥ 0. To our best knowledge, this problem has no explicit solution, and we use the R package `Rsolnp` (Ghalanos & Theussl 2015) to solve it numerically. In our estimation procedure, the averaging of matrix **M**_*l*_ over SNPs implies that higher-order terms of the allele frequency like *p*^2^ and *p*^3^ are simply estimated by the average of quadratic and cubic allele frequencies. These estimators are not unbiased (Weir 1996, Wang 2022), but can eventually be corrected for bias due to small sample size. One can thus correct for statistical sampling though this will not correct for genetic sampling (Weir 1996). No correction for statistical sampling was applied, given the large sample size used in our simulations below.

## 3 Simulations

We designed a simulation study for assessing the quality of the cls-based estimator for Jacquard coefficients and derived inbreeding and coancestry coefficients, and compared these with estimates obtained by em (Laporte et al. 2017). We simulated a pedigree with 20 unrelated founders, 10 males and 10 females, generating six non-overlapping generations totalling 111 individuals using the R-package `JGteach` (Goudet 2022). For more details on the pedigree-based simulation of genetic markers, see Goudet et al. (2018) and Leal et al. (2005). A picture of the simulated pedigree is shown in supplementary Figure S2. For each generation, females had a fertility rate of two, and fifty percent of the males were breeding. We generated 20,000 bi-allelic loci on a map of five Morgans. These settings allowed a considerable degree of relatedness to build up within a few generations. R (R Core Team 2023) instructions for generating the data are given in Appendix C. In order to assess the quality of the different estimators, we use the root-mean-squared-error (rmse). This requires the mathematical expectation of the various estimators of the Jacquard coefficients; in most cases this expectation is not available or hard to derive. We therefore use Jacquard coefficients calculated on realized ibd in the simulated pedigree as a proxy for this expectation, and calculate all rmse statistic with respect to these realized coefficients, which we call gold standard or just gold Jacquard coefficients. Likewise, in rmse calculations for derived coefficients (inbreeding, coancestry, etc.) we will also use gold values for these coefficients, which are obtained by applying the equations of Section 2 to the gold standard Jacquard coefficients. Figure 2 show scatterplots of the estimated Jacquard coefficients against their gold values for em and cls respectively. These plots are very noisy, revealing large errors in particular for estimates 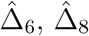 and 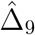, and show it is very hard to estimate the gold Jacquard coefficients reliably. Table 2 reports the RMSE for each coefficient for both methods. This shows the cls estimates have, over all, a lower rmse. The table confirms Δ_6_, Δ_8_ and Δ_9_ are most poorly estimated. In relatedness studies it is common practice to filter out low maf variants. Previous simulation work by Weir and Goudet (2017) has shown this leads to biased estimation of coancestry for the allele-sharing estimator of coancestry. We investigated the effect of MAF filtering by applying three MAF filters for both methods, and considering both the standard Jacquard-coefficient derived 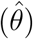 and the relative 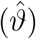 estimates of coancestry and inbreeding. The results in Table 2 show that maf filtering at one or five percent is detrimental for both the em and the cls algorithm, whereas leaving out monomorphic SNPS does not seem to affect the rmse. The negative effect of maf filtering is observed for both the standard parameters as well as relative coancestry and inbreeding. The estimates of the relative quantities have, in general, a slightly lower rmse. Consequently, the plots of our simulation results below used no maf filter and included all snps.

**Table 2:**
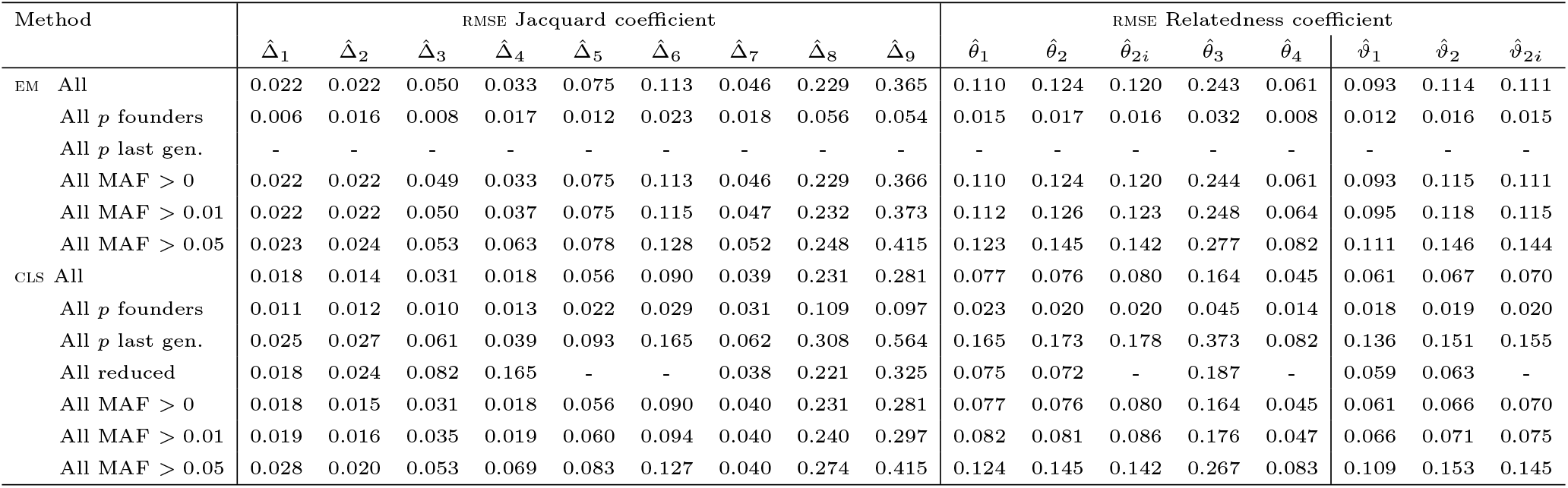
rmse of estimated relatedness parameters with respect to the gold Jacquard coefficients and gold relatedness parameters, for em and cls. The diagonals of 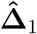 and 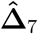 are not considered in the calculation of the rmse. All: using all individuals in the pedigree; All *p* founders: using all individuals and using founder allele frequencies. All *p* last gen.: using all individuals and allele frequencies of the last generation; All msf *>* 0: excluding monomorphic variants; All maf *>* 0.01 or *>* 0.05: using all individuals excluding variants with maf below the given threshold. All reduced: using all individuals and estimating the reduced system.

| Method |  | RMSE Jacquard coefficient |  |  |  |  |  |  |  |  | RMSE Relatedness coefficient |  |  |  |  |  |  |  |  |
| --- | --- | --- | --- | --- | --- | --- | --- | --- | --- | --- | --- | --- | --- | --- | --- | --- | --- | --- | --- |
| | | $\hat{\Delta}_1$ | $\hat{\Delta}_2$ | $\hat{\Delta}_3$ | $\hat{\Delta}_4$ | $\hat{\Delta}_5$ | $\hat{\Delta}_6$ | $\hat{\Delta}_7$ | $\hat{\Delta}_8$ | $\hat{\Delta}_9$ | $\hat{\theta}_1$ | $\hat{\theta}_2$ | $\hat{\theta}_{2i}$ | $\hat{\theta}_3$ | $\hat{\theta}_4$ | $\hat{\vartheta}_1$ | $\hat{\vartheta}_2$ | $\hat{\vartheta}_{2i}$ | |
| EM | All | 0.022 | 0.022 | 0.050 | 0.033 | 0.075 | 0.113 | 0.046 | 0.229 | 0.365 | 0.110 | 0.124 | 0.120 | 0.243 | 0.061 | 0.093 | 0.114 | 0.111 |  |
| | All $p$ founders | 0.006 | 0.016 | 0.008 | 0.017 | 0.012 | 0.023 | 0.018 | 0.056 | 0.054 | 0.015 | 0.017 | 0.016 | 0.032 | 0.008 | 0.012 | 0.016 | 0.015 | |
| | All $p$ last gen. | - | - | - | - | - | - | - | - | - | - | - | - | - | - | - | - | - | |
|  | All MAF > 0 | 0.022 | 0.022 | 0.049 | 0.033 | 0.075 | 0.113 | 0.046 | 0.229 | 0.366 | 0.110 | 0.124 | 0.120 | 0.244 | 0.061 | 0.093 | 0.115 | 0.111 |  |
|  | All MAF > 0.01 | 0.022 | 0.022 | 0.050 | 0.037 | 0.075 | 0.115 | 0.047 | 0.232 | 0.373 | 0.112 | 0.126 | 0.123 | 0.248 | 0.064 | 0.095 | 0.118 | 0.115 |  |
|  | All MAF > 0.05 | 0.023 | 0.024 | 0.053 | 0.063 | 0.078 | 0.128 | 0.052 | 0.248 | 0.415 | 0.123 | 0.145 | 0.142 | 0.277 | 0.082 | 0.111 | 0.146 | 0.144 |  |
| CLS | All | 0.018 | 0.014 | 0.031 | 0.018 | 0.056 | 0.090 | 0.039 | 0.231 | 0.281 | 0.077 | 0.076 | 0.080 | 0.164 | 0.045 | 0.061 | 0.067 | 0.070 |  |
| | All $p$ founders | 0.011 | 0.012 | 0.010 | 0.013 | 0.022 | 0.029 | 0.031 | 0.109 | 0.097 | 0.023 | 0.020 | 0.020 | 0.045 | 0.014 | 0.018 | 0.019 | 0.020 | |
| | All $p$ last gen. | 0.025 | 0.027 | 0.061 | 0.039 | 0.093 | 0.165 | 0.062 | 0.308 | 0.564 | 0.165 | 0.173 | 0.178 | 0.373 | 0.082 | 0.136 | 0.151 | 0.155 | |
|  | All reduced | 0.018 | 0.024 | 0.082 | 0.165 | - | - | 0.038 | 0.221 | 0.325 | 0.075 | 0.072 | - | 0.187 | - | 0.059 | 0.063 | - |  |
|  | All MAF > 0 | 0.018 | 0.015 | 0.031 | 0.018 | 0.056 | 0.090 | 0.040 | 0.231 | 0.281 | 0.077 | 0.076 | 0.080 | 0.164 | 0.045 | 0.061 | 0.066 | 0.070 |  |
|  | All MAF > 0.01 | 0.019 | 0.016 | 0.035 | 0.019 | 0.060 | 0.094 | 0.040 | 0.240 | 0.297 | 0.082 | 0.081 | 0.086 | 0.176 | 0.047 | 0.066 | 0.071 | 0.075 |  |
|  | All MAF > 0.05 | 0.028 | 0.020 | 0.053 | 0.069 | 0.083 | 0.127 | 0.040 | 0.274 | 0.415 | 0.124 | 0.145 | 0.142 | 0.267 | 0.083 | 0.109 | 0.153 | 0.145 |  |

**Figure 2.**
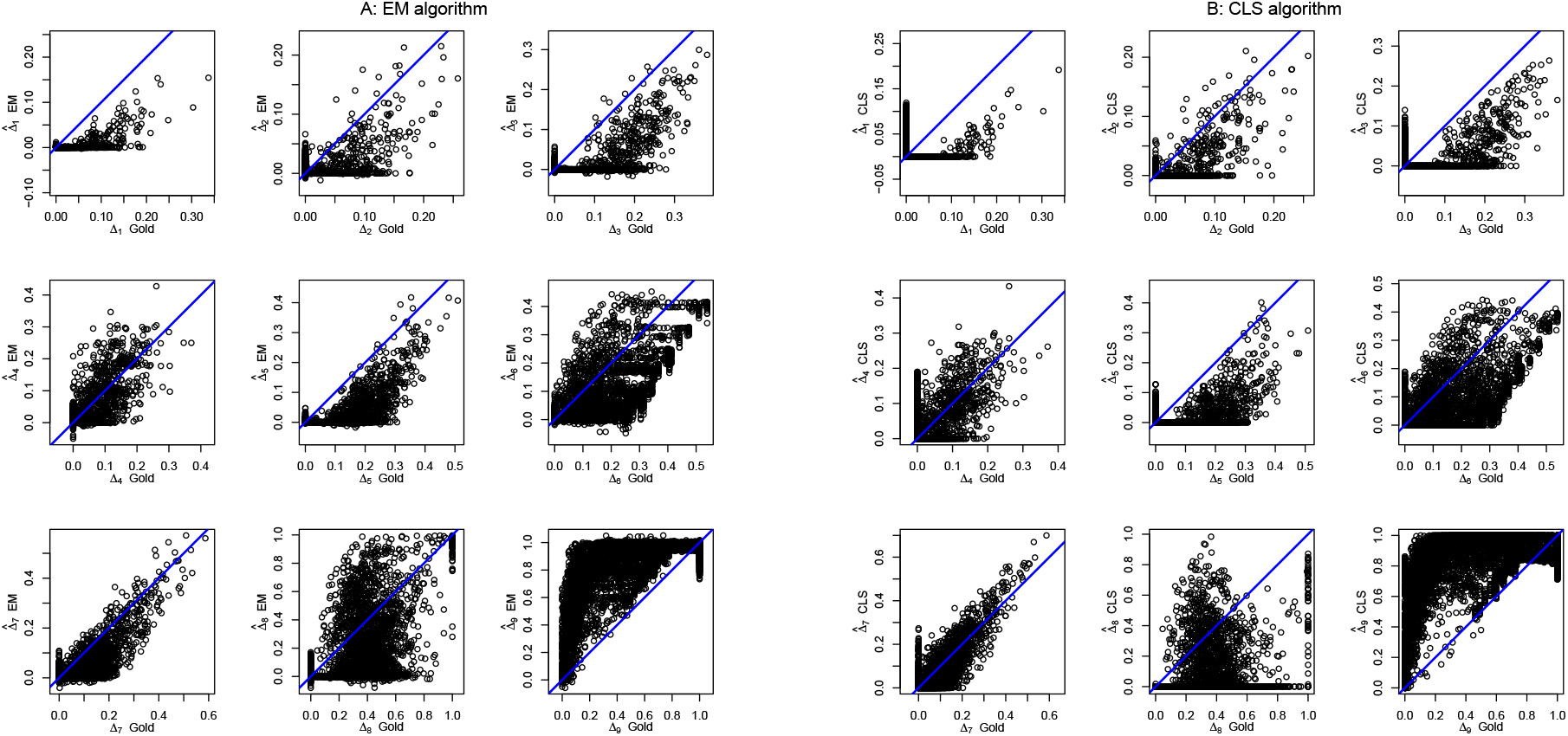
Estimation of gold Jacquard coefficients by em (panel A) and cls (panel B).

The poor estimation results are probably in part explained by the fact the coefficients are not identified in the bi-allelic case; it is of more interest to focus on derived quantities that are identifiable: coancestry, inbreeding and other relatedness parameters (Csűrös 2014). Figure 3 shows scatterplots of the estimated coancestry and inbreeding coefficients against their gold values for em and cls respectively, and the corresponding rmses are given in the last eight columns of Table 2. Figure 3 and Table 2 both show that cls estimates have, in general, less variation and come closer the the *y* = *x* line. However, both estimators substantially underestimate coancestry, inbreeding and the probability that least one ibd pair out of three (*θ*_3_).

**Figure 3.**
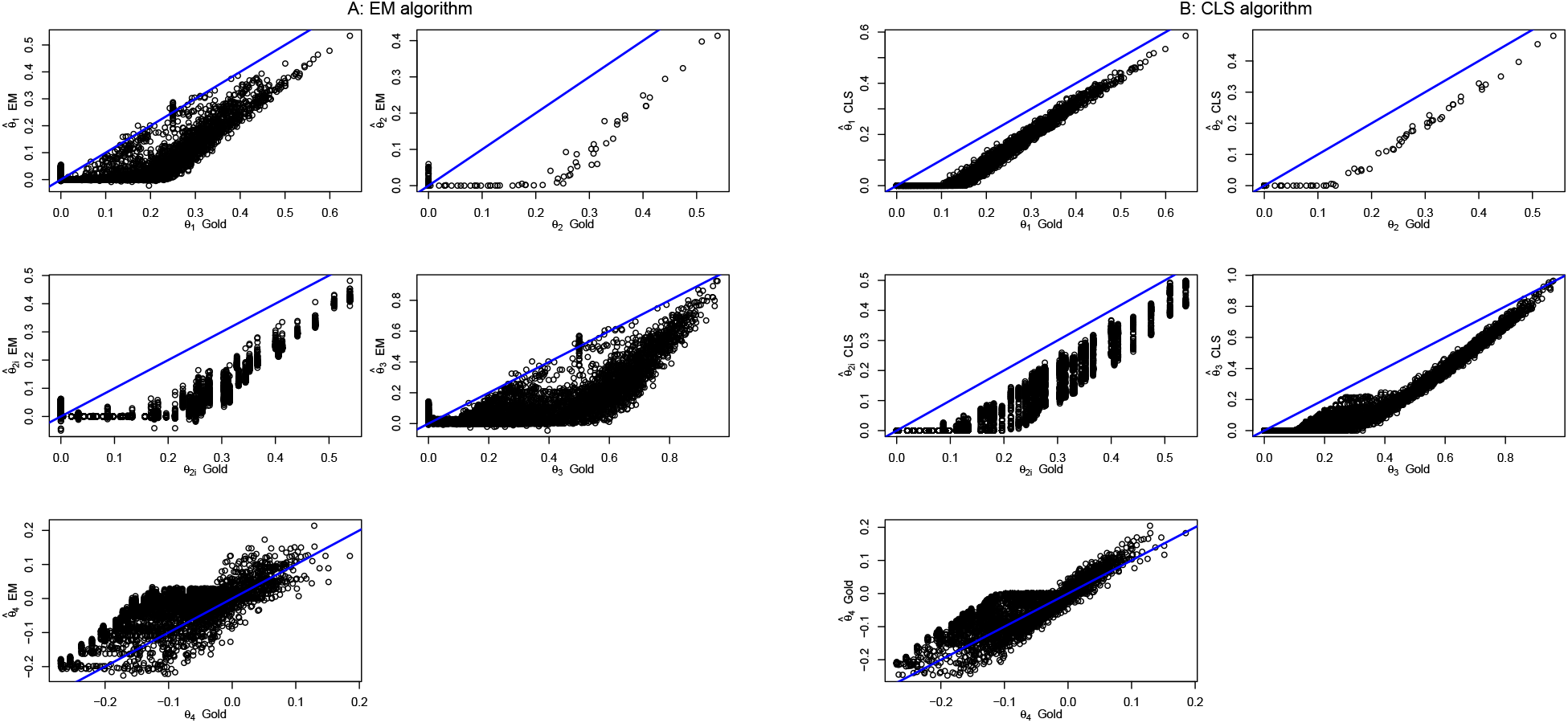
Estimation of relatedness parameters by em (panel A) and cls (panel B). For inbreeding, both the individual inbreeding coefficients 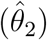 and the corresponding pairwise estimates 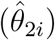 are shown.

Weir and Goudet (2017, Tables 1 and 3) proposed unbiased allele-sharing estimators for the compound quantities of *relative* coancestry and *relative* inbreeding, which are obtained as follows:

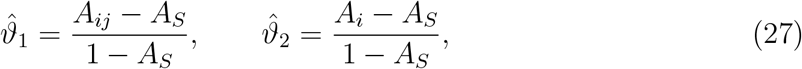

where *A*_*ij*_ and *A*_*i*_ are allele-sharing statistics, with *A*_*ij*_ the proportion of alleles carried by individuals *i* and *j* that are identical in state (ibs), *A*_*i*_ the proportion of loci for which individual *i* is homozygous, and *A*_*S*_ the average of *A*_*ij*_ over all pairs of distinct individuals 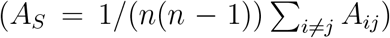. We suggest to convert the em and cls estimators for coancestry 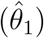 and inbreeding 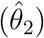, which are obtained from the estimated Jacquard coefficients, into estimators of the relative compound quantities, by using a transformation inspired by Eqs. (22) and (27):

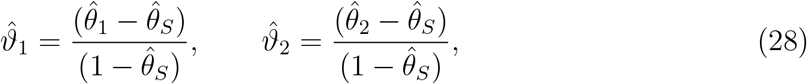

where 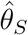 is the sample average of all pairwise coancestry estimates 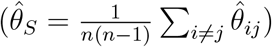.

We note that the allele-sharing estimators directly estimate the relative quantities of interest, whereas Eq. (28) modifies pre-existing estimates of *θ*_1_ and *θ*_2_. The resulting estimators may not be unbiased, though the simulations suggest the relative parameters are better estimated (see the last three columns of Table 2.) Moreover, the gold values of *θ*_1_ and *θ*_2_ are clearly underestimated by both algorithms (see Figure 3); this improves if Eq. (28) is used to estimate the relative gold values (see Figure 4). We note the sample average of the relative coancestry estimator is zero; in general this will not be the case for the relative inbreeding estimator. We also note the relative estimators amount to a linear rescaling of their original em and cls counterparts; consequently any estimator of coancestry will have the same correlation with 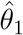 and 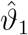; likewise for estimators of inbreeding and 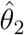 and 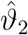. Figure 4 shows the estimation of the compound relative parameters is more successful for both the em and the cls algorithm, and leads to improved approximation of the gold values, as also witnessed by the RMSE statistics in Table 2. We note that gold values for inbreeding are constant across all pairs for a given individual, though the corresponding cls and em estimates fluctuate, since not all pairs converge to the same value.

**Figure 4.**
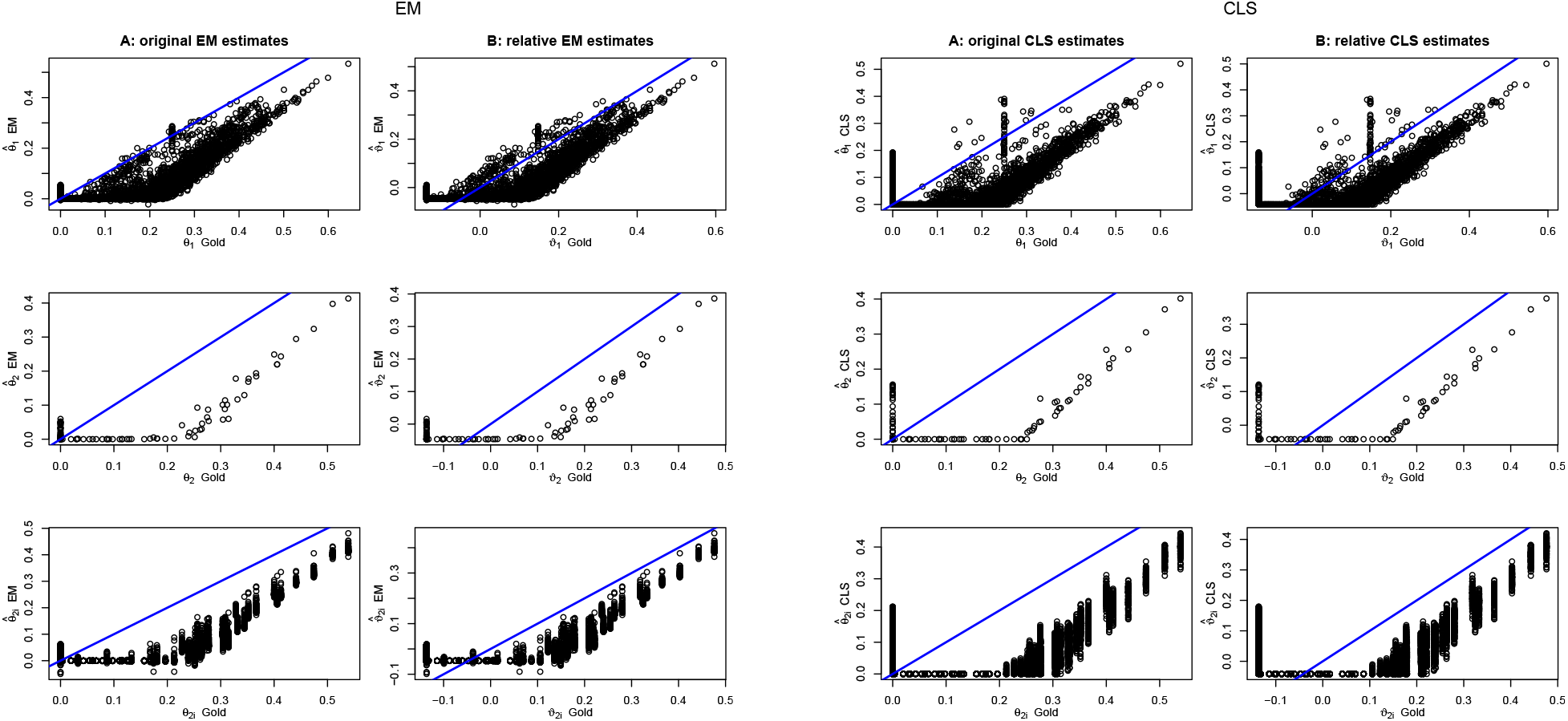
Estimation of gold coancestry and inbreeding coefficients with em and cls estimators. The left column (A:) of each panel shows the original gold values (*θ*_*i*_) against the estimates 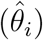 of the algorithm. The right column (B:) shows the *relative* gold values (*ϑ*_*i*_) against their *relative* estimates 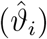. The first row shows coancestry, the second row individual inbreeding coefficients as estimated by 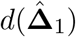, the last row shows all pairwise estimates for the inbreeding coefficient of a given individual obtained according to Eq. (14).

The quality of the different estimators depends on the sample allele frequencies and pairs of individuals that are used for comparison. In a simulation, for the estimation of the allele frequencies one can use founders, last-generation individuals or the full pedigree (as in Figures 2 and 3). When the estimation is carried out using only the last-generation individuals for estimating allele frequencies, rmse statistics generally deteriorate for CLS, whereas it was mostly impossible to obtain em estimates in most cases. This is a likely consequence of both a decrease in sample size by 87% and a considerable increase in the percentage of monomorphic snps (72% in the last generation, versus 11% in the founder generation). When founder allele frequencies are used for estimation, the fit, considering the full pedigree, improves considerably (see Table 2 and Figure 5). It reduces the rmse for all Jacquard estimates, and for 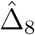 and 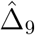 in particular, for both the em and CLS algorithm (see Table 2), and consequently also gives a lower rmse for the derived relatedness coefficients. With founder allele frequencies, rmse statistics for the em algorithm are in general, slightly lower than those of the cls algorithm, suggesting the allele frequencies are particularly crucial to the em algorithm. The best estimation results for coancestry and inbreeding are obtained by using the em algorithm with founder allele frequencies, and estimating the relative quantities that account for average coancestry. We also fitted the reduced bi-allelic system. Supplementary Figure S3 shows the estimates for the seven reduced Jacquard coefficients and coancestry, inbreeding and 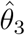 obtained by fitting the reduced bi-allelic system. Despite fitting two parameters less, the rmse statistics obtained for coancestry, inbreeding and *θ*_3_ are almost the same as obtained by fitting the nine parameter condensed system.

**Figure 5.**
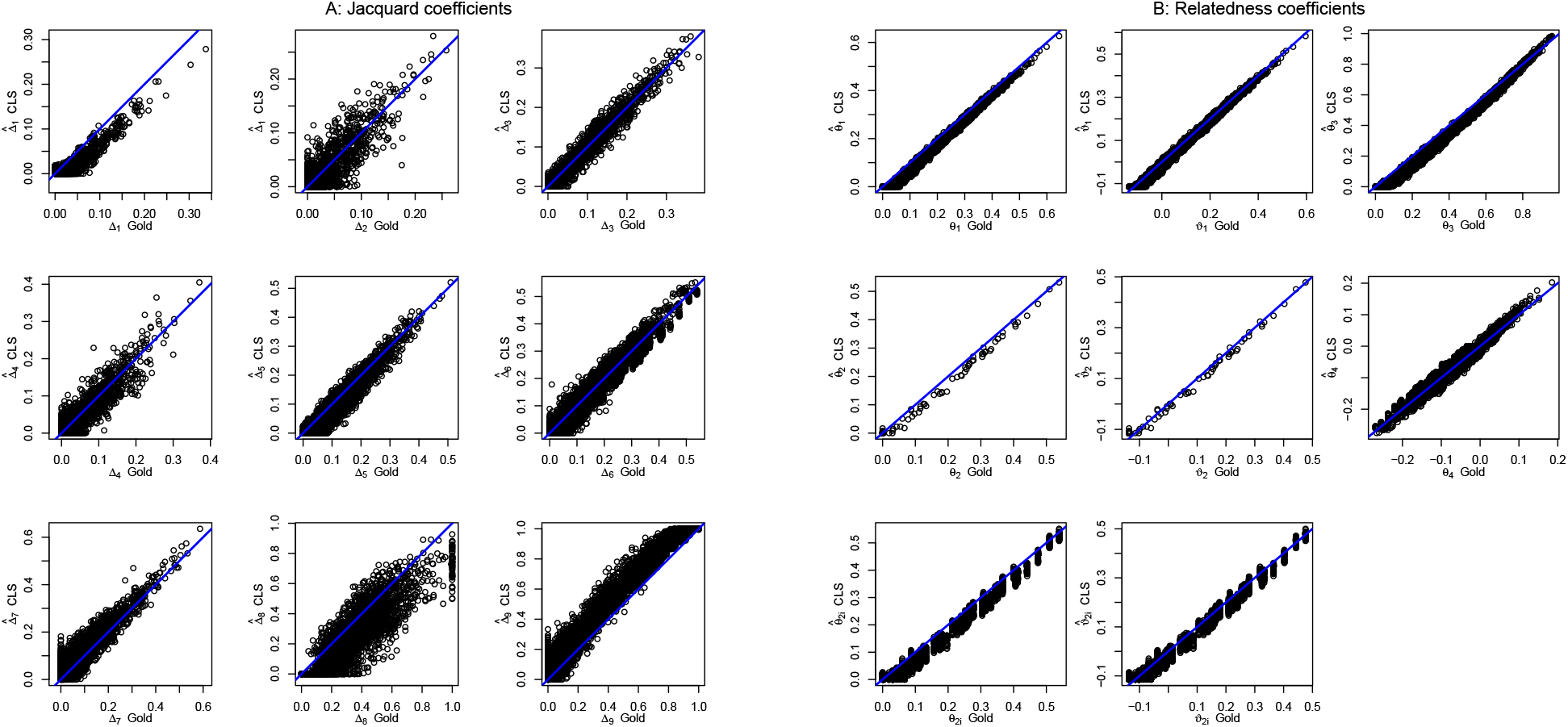
Estimation of gold Jacquard and relatedness coefficients by CLS with founder allele frequencies. For inbreeding, both individual estimates (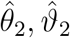, second row) and pairwise estimates (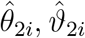 third row) are shown.

We explored the computational cost of scaling up both algorithms by using increasing numbers of SNPs, up to a million. For the em algorithm, we found estimation of the full set of Jacquard coefficient to be infeasible for larger numbers of SNPs. Figure 6 shows the CPU time spent for a sample of size 109. For both algorithms the CPU time increases, as expected, linearly with the number of polymorphisms. Figure 6 shows that the CLS algorithm (with tolerance parameter 1E-8) is much faster than the em algorithm (used with convergence precision 1E-3). The calculation of the Jacquard coefficients for one million SNPs required 48.86 hours for the em algorithm, whereas this takes only 0.36 hours for the CLS algorithm, where, for the sake of comparison, we used a single core.

**Figure 6.**
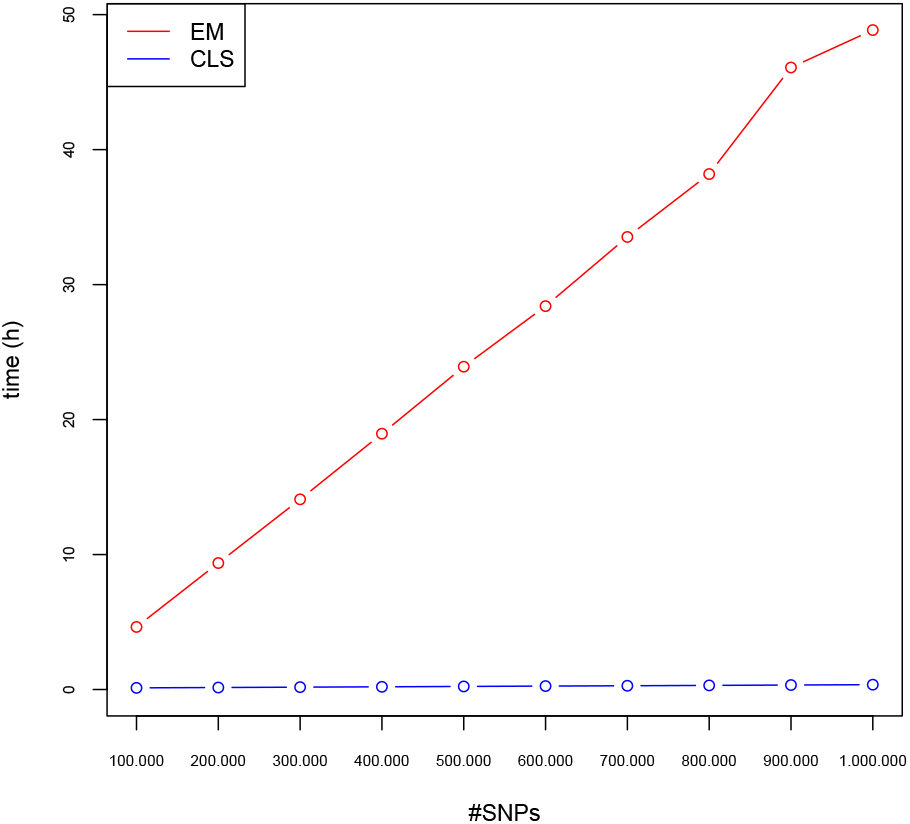
Computational cost (in hours) for estimation of the Jacquard coefficients as a function of the number of bi-allelic SNPs and algorithm used (em or cls).

## 4 Discussion

Considerable research effort has been dedicated to the estimation of relatedness parameters such as kinship and inbreeding coefficients. Hitherto, there is little work on the estimation of the full set of Jacquard’s genetic identity coefficients with the use of molecular marker data. For bi-allelic genetic variants, at first sight there may seem to be little point in reporting the full set because they are not identified (see Figure S1). Nevertheless, reporting the full set of coefficients is ultimately informative for it will always permit the calculation of any identifiable derived relatedness parameter. Maximum likelihood estimation by means of the em algorithm (Laporte et al. 2017) is computationally very expensive and not feasible on a genomewide scale. The proposed cls approach is seen to provide comparable estimates of the Jacquard coefficients and derived quantities, and simulations suggest these have smaller rmse when the founder allele frequencies are unknown. The likelihood approach is probabilistic and multiplies over presumably independent loci, whereas such independence is known not to hold for markers on the same chromosome that are close. The proposed cls approach is purely based on least-squares minimization and makes no implicit assumptions about ld.

Our simulations show that the estimation of Jacquard and relatedness coefficients works best with founder allele frequencies. Founder allele frequencies will usually be available in breeding programs, but remain unknown in many other empirical settings, where estimation of allele frequencies will typically be based on all available individuals.

The cls approach proposed in this article is flexible, and can be further extended for variants with multiple alleles, such as microsatellites. In that case, the identification problem of the Jacquard coefficients is resolved. It is also easily adapted for the classical estimation, under the assumption of no inbreeding, of the Cotterman coefficients. In order to do so, one should just carry out the minimization while restricting the first six Jacquard coefficients to be zero and additionally, imposing Thompson’s (1976) condition for the absence of inbreeding. A common sense data-analytic strategy is to first estimate the full set of Jacquard coefficients without any inbreeding constraint, but to impose Thompson’s constraint in second instance in case no obvious evidence for inbreeding is found. Interestingly, if pairs are known (or believed) to be unrelated, this condition may be imposed by restricting all related states (Δ_1_, Δ_3_, Δ_5_, Δ_7_ and Δ_8_) to be zero and estimating only Δ_9_ and the remaining inbred states. Indeed, any subset of the Jacquard coefficients may be set to zero as suggested by the results of a first exploratory analysis, and to the benefit of reducing the identifiability problem.

Both the em and cls algorithms adhere to a strict probabilistic interpretation of Jacquard’s coefficients, their estimates can not be negative and consequently inbreeding and coancestry estimates can neither be negative. The simulations suggest improved approximation of gold inbreeding and coancestry may be possible if negative values would be admitted (see Figures 3 and 4), though this is clearly less compelling if better estimates of the allele probabilities are available, as is the case with founder allele frequencies (see Figure 5). Also, the flooring of coancestry estimates at zero pulls estimates of average coancestry towards zero and impacts the correction for average coancestry. em and cls algorithms could be further developed towards explicitly estimating the relative quantities of interest, and possibly lifting the non-negativity constraint.

There are some considerations that may be helpful to reduce the computational burden. When the pairs of individuals of interest are known in advance, the expensive calculation of all relatedness statistics for all pairs can be avoided. To obtain the statistics of interest, one only needs to calculate the allele frequencies, and subset the calculations of the relatedness statistics to the genotype data of the pairs of interest only. This applies to both the cls and the em algorithm, as both operate in a pairwise manner. If the estimation of inbreeding is of main interest, for the CLS approach the pairwise calculations can be greatly reduced, because in that case only *n* estimates of the first Jacquard coefficient of an individual are needed instead of the usual 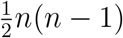 pairs. Many genetic studies filter genetic variants by their MAF, with MAF ≤ 0.05 being a commonly used exclusion criterion. Given the typically skewed distribution of the MAF in empirical studies, such filtering implies the exclusion of huge amounts of polymorphisms, and can so reduce computational cost. However, previous simulation work of Weir and Goudet (2017) has shown that MAF filtering introduces bias in the estimation of (pedigree-based) coancestry, whereas our simulations in Table 2 show increased rmse for all Jacquard coefficients and derived quantities.

Estimation of the Jacquard coefficients with either em or cls relies on numerical optimization for which global convergence is not always guaranteed. Proper convergence can be investigated by modifying the tolerance criterion for convergence and by choosing different initial points. If maximum likelihood estimation is preferred, the CLS estimates can be used as initial points for the em algorithm, to the benefit of the convergence of the latter.

Both the em algorithm and the proposed CLS approach relies on adequate estimates of the allele frequencies, for which sample allele frequencies are typically used, obtained from either the full data set, or, if possible, from the founder generation only. Reliance on sample allele frequencies is the current approach in relatedness research, as most kinship and inbreeding coefficient estimators do require allele frequency estimates. Alternatively, allele sharing estimators that avoid the use of sample allele frequencies have recently been developed (Goudet et al. 2018, Weir & Goudet 2017). The latter estimators do not provide the full set of Jacquard coefficients, but for estimating coancestry and inbreeding they do not rely on iterative algorithms and are computationally very cheap.

## 5 Software

Estimates of Jacquard’s coefficients with the em algorithm were obtained with the R package `Relatedness` (Laporte & Mary-Huard 2017, Laporte et al. 2017). Simulated pedigrees used in this article were generated with R package `JGTeach` (Goudet 2022). We developed R package `Jacquard` which implements estimation of Jacquard’s coefficients and derived quantities by constrained least squares, relying on the optimization functions of R package `Rsolnp` (Ghalanos & Theussl 2015).

## 6 Acknowledgements

This work was supported by the Spanish Ministry of Science and Innovation and the European Regional Development Fund under grant PID2021-125380OB-I00 (MCIN/AEI/FEDER); and the National Institutes of Health under Grant GM075091.

The authors report there are no competing interests to declare.

## 7 Supplementary Material

## Appendix A: Identifiability of Jacquard’s coefficients

We illustrate the identifiability problem of the Jacquard coefficients with an example, using the vector of joint genotype probabilities, in the order of Table 1, (0.356, 0.012, 0.038, 0.038, 0.141, 0.033, 0.034, 0.175, 0.173) with associated allele frequency *p* = 0.26. The effective constraints for (*ξ, η*) are shown as lines in Figure S1A. Any point within the central polygon can map the negative Jacquard coefficients into non-negative probabilities. A feasible point (*ξ, η*) that produces non-negative Jacquard coefficients can in this case be found by averaging the upper and lower limit of the active constraints, as shown by the single dot in the Figure S1A. By exploring a grid of (*ξ, η*) values over the central polygon, sets of non-negative Jacquard coefficients are obtained whose range of variation is shown in Figure S1B. This shows a certain variation in these coefficients (Δ_8_ in particular), defining sets that are compatible with the same linear system of equations. When the kinship, inbreeding and other relatedness coefficients are calculated from the feasible Jacquard coefficients, Figure S1C is obtained, illustrating that these relatedness coefficients are virtually constant and thus identified.

**Figure S1:**
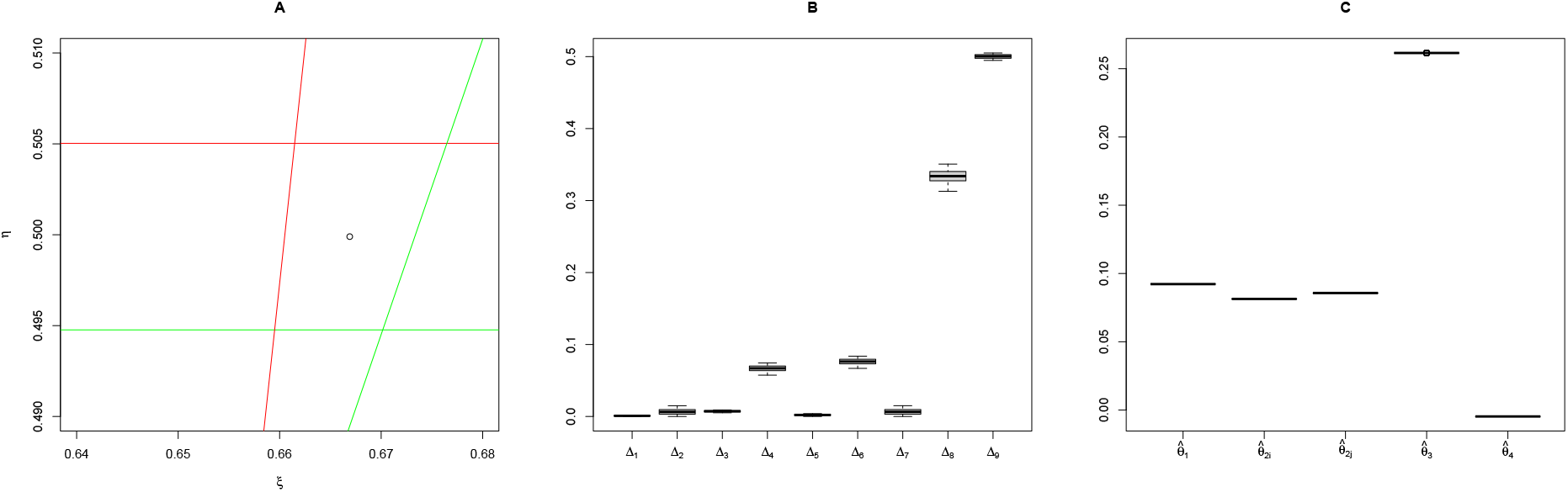
A: Inequalities determining (*ξ, η*) that map Jacquard coefficients to probabilities. B: Variation in the Jacquard coefficients obtained by varying (*ξ, η*) over the polygon in panel A. C: Kinship, inbreeding and other relatedness coefficients obtained from the Jacquard coefficients in panel B.

For a bi-allelic system, the Jacquard coefficients could be made identifiable, but this would require the imposition of additional constraints on the coefficients. Alternatively, with a multiallelic system, matrix **M** with the conditional joint genotype probabilities, as laid out by Anderson and Weir (2007) or Wang (2022), is generally full rank and in that case the Jacquard coefficients are identifiable.

## Appendix B: Reduction of the number of Jacquard coefficients

We will use Δ_35_ to represent the sum of Δ_3_ and Δ_5_; likewise Δ_46_ = Δ_4_+Δ_6_. We call the new set of seven coefficients the reduced Jacquard coefficients. Importantly, the joining of these columns creates additional linear dependence between the rows of Table 1, where three pairs of rows ((2,4), (3,7) and (6,8)) become identical. We next sum the two rows of each pair, which results in the coefficient matrix of a *biallelic reduced system* given in Table S1. This system retains a linear constraint on the columns, and can again be estimated by constrained least squares with the partly combined Jacquard coefficients having weighted sum one. Again importantly, the joining of the rows precisely eliminates the ordering of the genotype pairs; i.e. the probabilities of (0/0,0/1) and (0/1,0/0) are now summed. As a consequence, the new coefficient matrix **M**^∗^ is now of dimension 6 *×* 7 and will generally have rank five due to the remaining linear restriction on the sum of the columns; in the exceptional case *p* = *q* = 0.5 it will have rank four. This fusion leaves the theoretical relatedness parameters *θ*_1_ and *θ*_3_ unaltered. The inbreeding coefficients *θ*_2*i*_ and *θ*_2*j*_ can no longer be independently estimated, though we can now estimate the probability that either *i* or *j* is inbred, and by averaging these probabilities over pairs, an individual inbreeding coefficient can be estimated. Parameter *θ*_4_ (Eq. (6)) can no longer be estimated.

**Table S1:**
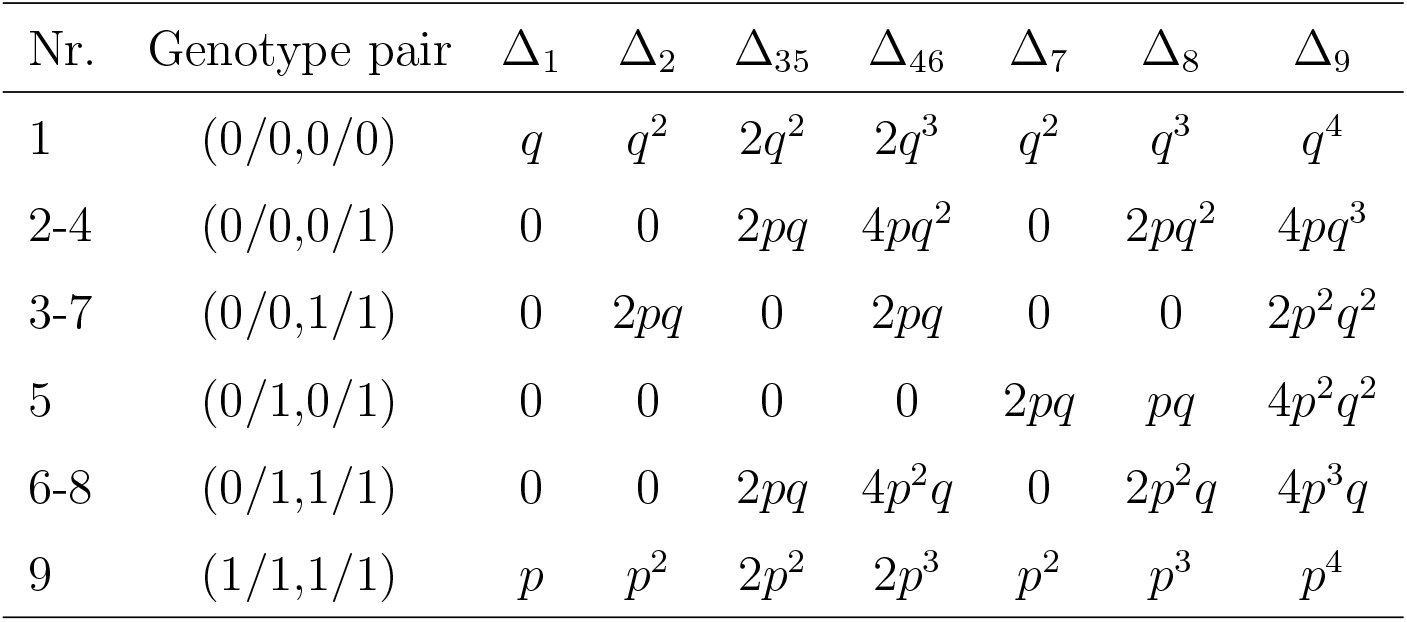
The biallelic reduced condensed system, consisting of unordered joint genotype probabilities for given ibd patterns and allele frequencies.

| Nr. | Genotype pair | $\Delta_1$ | $\Delta_2$ | $\Delta_{35}$ | $\Delta_{46}$ | $\Delta_7$ | $\Delta_8$ | $\Delta_9$ |
| --- | --- | --- | --- | --- | --- | --- | --- | --- |
| 1 | (0/0,0/0) | $q$ | $q^2$ | $2q^2$ | $2q^3$ | $q^2$ | $q^3$ | $q^4$ |
| 2-4 | (0/0,0/1) | 0 | 0 | $2pq$ | $4pq^2$ | 0 | $2pq^2$ | $4pq^3$ |
| 3-7 | (0/0,1/1) | 0 | $2pq$ | 0 | $2pq$ | 0 | 0 | $2p^2q^2$ |
| 5 | (0/1,0/1) | 0 | 0 | 0 | 0 | $2pq$ | $pq$ | $4p^2q^2$ |
| 6-8 | (0/1,1/1) | 0 | 0 | $2pq$ | $4p^2q$ | 0 | $2p^2q$ | $4p^3q$ |
| 9 | (1/1,1/1) | $p$ | $p^2$ | $2p^2$ | $2p^3$ | $p^2$ | $p^3$ | $p^4$ |

For the reduced condensed system, the previously developed matrix equations largely apply with some modifications, in particular

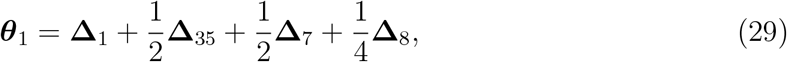

and a matrix of inbreeding coefficients is obtained as

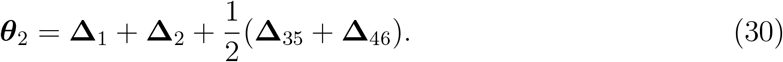

The previous equations (12), (13) and (17) still apply, and for the newly defined inbreeding coefficients we have that

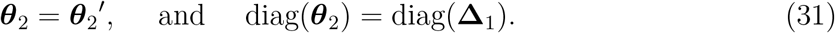

The probability of at least one pair of ibd alleles among three randomly selected alleles is now given by

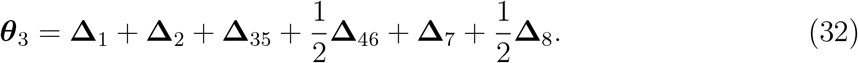

## Appendix C: R instructions for the simulation

~~~
library(JGTeach) library(hierfstat)
set.seed(1234)
X.ped <- buildped.2sexes(founders.m=10,founders.f=10,fert=2,
 death.rate=1,breed.prop=0.5,n.tstep=6)
set.seed(123)
PG <- drop.along.ped(X.ped,nloc=20000,maplength=5) X.gen <- IBD2dos(PG,20)
~~~

## Appendix D: Supplemental figures for the simulations

**Figure S2:**
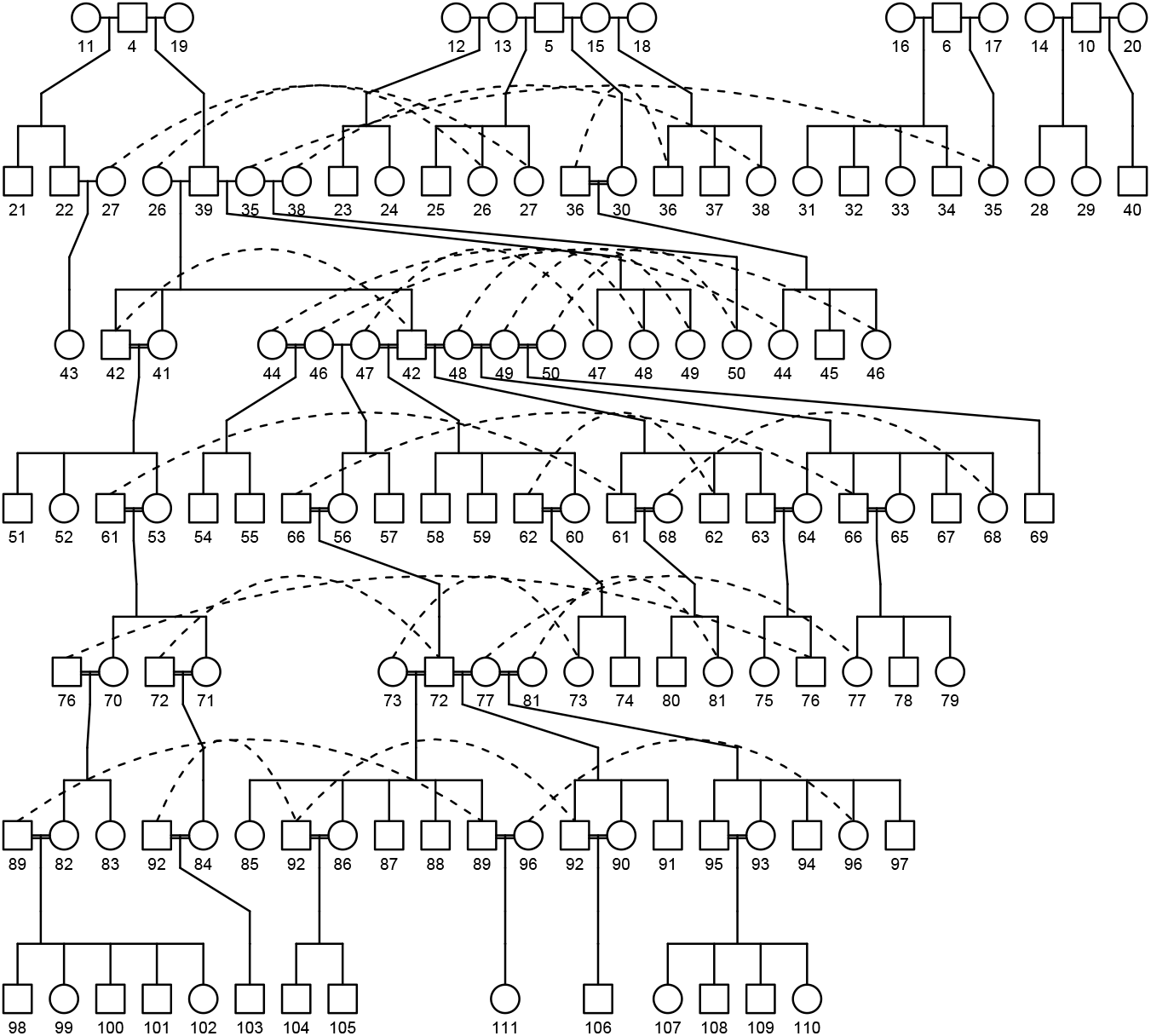
Pedigree simulated by the `JGteach` R-package spanning seven generations (founders included).

**Figure S3:**
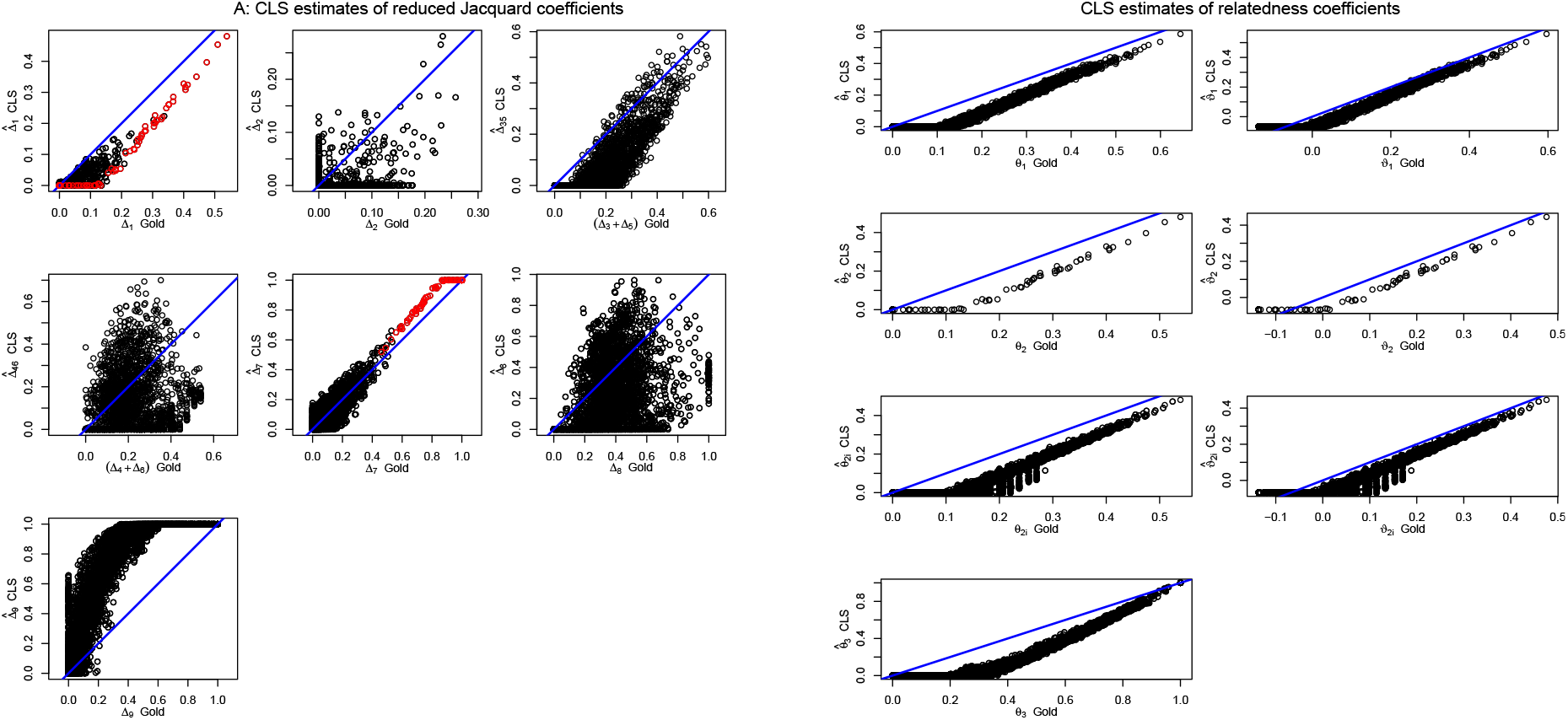
Estimation of reduced Jacquard coefficients. A: Seven reduced Jacquard coefficients versus their gold values. Red dots indicate diagonal elements of **Δ**_1_ and **Δ**_7_. B: Relatedness coefficients. The left column represents the original IBD based estimates, the right column the relative estimates.

## Notes

### Competing Interest Statement

The authors have declared no competing interest.

